# A dual DNA/RNA-binding factor regulates co-transcriptional splicing through target RNA interaction and modulates splicing factor dynamics

**DOI:** 10.1101/2024.01.11.575216

**Authors:** Mukulika Ray, Julia Zaborowsky, Pranav Mahableshwarkar, Smriti Vaidyanathan, Jasmine Shum, Annie Huang, Renjith Viswanathan, Isabelle Pilo, Victoria Chen, Kaitlyn Cortez, Ashley M. Conard, Szu-Huan Wang, Victoria Johnson, Noah Wake, Alexander E. Conicella, Ryan Puterbaugh, Nicolas L. Fawzi, Erica Larschan

**Author notes:** Co-first Authors (Mukulika Ray, Julia Zaborowsky).

## Abstract

How RNA splicing events are targeted to the correct genomic locations in specific cellular contexts to generate context-specific transcript diversity and prevent deleterious cryptic splicing remains very poorly understood. We show that a functionally conserved GA-rich DNA-binding transcription factor (TF), CLAMP, targets distinct RNA molecules in male and female cells to precisely regulate sex-specific splicing events through physical and functional interactions with RNA and RNA-binding proteins (RBPs). The prion-like domain of CLAMP (PrLD) and a stem-loop region in the target RNA are important for CLAMP-RNA interaction. Moreover, the CLAMP PrLD domain regulates sex-specific splicing by modulating the dynamics of an hnRNPA2/B1 family protein that regulates alternative splicing. Thus, we demonstrate that a TF targets co-transcriptional splicing to the correct genomic locations by directly linking DNA binding sites to RNA targets and modulating the dynamics of RBP partners that drive alternative splicing.

## Introduction

Mechanisms that drive precise co-transcriptional alternative splicing at thousands of genomic loci are critical for establishing sex- and cell-type-specific transcriptome diversity and preventing deleterious cryptic splicing events that often occur in disease contexts^1^. Chromatin-bound transcription factors (TFs) and RNA-binding proteins (RBPs) co-transcriptionally regulate alternative splicing^2–5^. Most TFs bind DNA and RNA^6–8^ and interact with diverse RBPs^9,10^. However, the mechanisms by which TFs and RBPs coordinate transcription and alternative splicing at specific genomic locations in a particular cell type to drive specific alternative splicing events and prevent deleterious cryptic splicing remain poorly understood. Also, how interactions between TFs, RNA, and RBPs generate the specific alternatively spliced transcripts essential for making developmental decisions remains unknown. We hypothesized that a subset of TFs are critical for targeting the correct co-transcriptional splicing events to specific genomic locations due to their unique ability to bind chromatin, RNA, and spliceosomal RBPs at intron-exon boundaries.

We test this hypothesis by defining the role of the GA-binding TF CLAMP (chromatin-linked adapter for MSL Proteins) in targeting sex-specific splicing to the correct genomic locations on chromatin in *Drosophila* because we can simultaneously genetically manipulate the system, conduct high-throughput genomic experiments *in vivo,* and performing live imaging experiments in living tissue. CLAMP has several properties that suggest that it could target sex-specific alternative splicing events to chromatin: 1) CLAMP binds to specific GA-enriched DNA motifs via its mapped DNA binding domain, and its binding sites differ between males and females^11–14^; 2) CLAMP is a pioneer TF that regulates sex-specific splicing in embryos at genes where it does not regulate transcription^15^; 3) CLAMP is enriched at the intronic regions of CLAMP-dependent sex-specifically spliced genes^12,15^; 4) CLAMP is associated with RBPs that are spliceosome components^10^.

Here, we combine diverse *in vivo* and *in vitro* approaches to define a new mechanism by which a TF can regulate co-transcriptional splicing. We define direct CLAMP-RNA interactions and compare them with CLAMP-DNA interactions to identify associations between TF-DNA and TF-RNA binding at genes regulated by CLAMP in sex-specific alternative splicing. We have identified essential protein and RNA domains that promote direct CLAMP-RNA interactions using NMR (Nuclear Magnetic Resonance) spectroscopy and RNA-protein EMSAs (Electrophoretic Mobility Shift Assays). Furthermore, we determined that CLAMP directly regulates the biophysical properties of Hrp38, its RBP interaction partner, which is an hnRNPA2/B1 homolog known to regulate alternative splicing as part of nuclear splicing condensates. Thus, overall, our data support a model in which transcription factors with both DNA and RNA binding capacity can target unique RNA molecules to specific genomic locations and alter the biophysical properties of RBP partner proteins to regulate context-specific alternative splicing.

## Results

### CLAMP binds RNA on chromatin

Because CLAMP is a DNA-binding protein enriched at the intronic regions of CLAMP-dependent sex-specifically spliced genes, which interacts with multiple splicing factors^12,15^, we hypothesized that CLAMP functions as a regulator of co-transcriptional splicing. To test this hypothesis, we first investigated whether CLAMP regulates sex-specific splicing by directly interacting with target RNAs. Therefore, we defined RNA species interacting with CLAMP at high resolution across different cellular compartments: the nucleoplasm vs. chromatin. If we identified CLAMP-RNA interactions in the chromatin fraction involving RNAs that require CLAMP for their alternative splicing, this would support a model in which CLAMP targets co-transcriptional splicing to specific genomic loci. To further define a potential co-transcriptional function for CLAMP, we wanted to identify the spatial distribution of CLAMP RNA binding with respect to its DNA binding across the genome by comparing CLAMP-RNA binding sites with CLAMP DNA binding sites.

To achieve these objectives, we performed fractionation iCLIP (**i**ndividual-nucleotide resolution **C**ross**L**inking and **I**mmuno**P**recipitation) to identify CLAMP RNA targets using anti-CLAMP antibody in chromatin fractions (ChF) and nucleoplasmic fractions (NF) of male (S2) and female (Kc) *Drosophila* cell lines (Fig 1A). Kc and S2 cells are well-established cell lines for studying sex-specific processes such as dosage compensation^16,17^.

**Figure 1.**
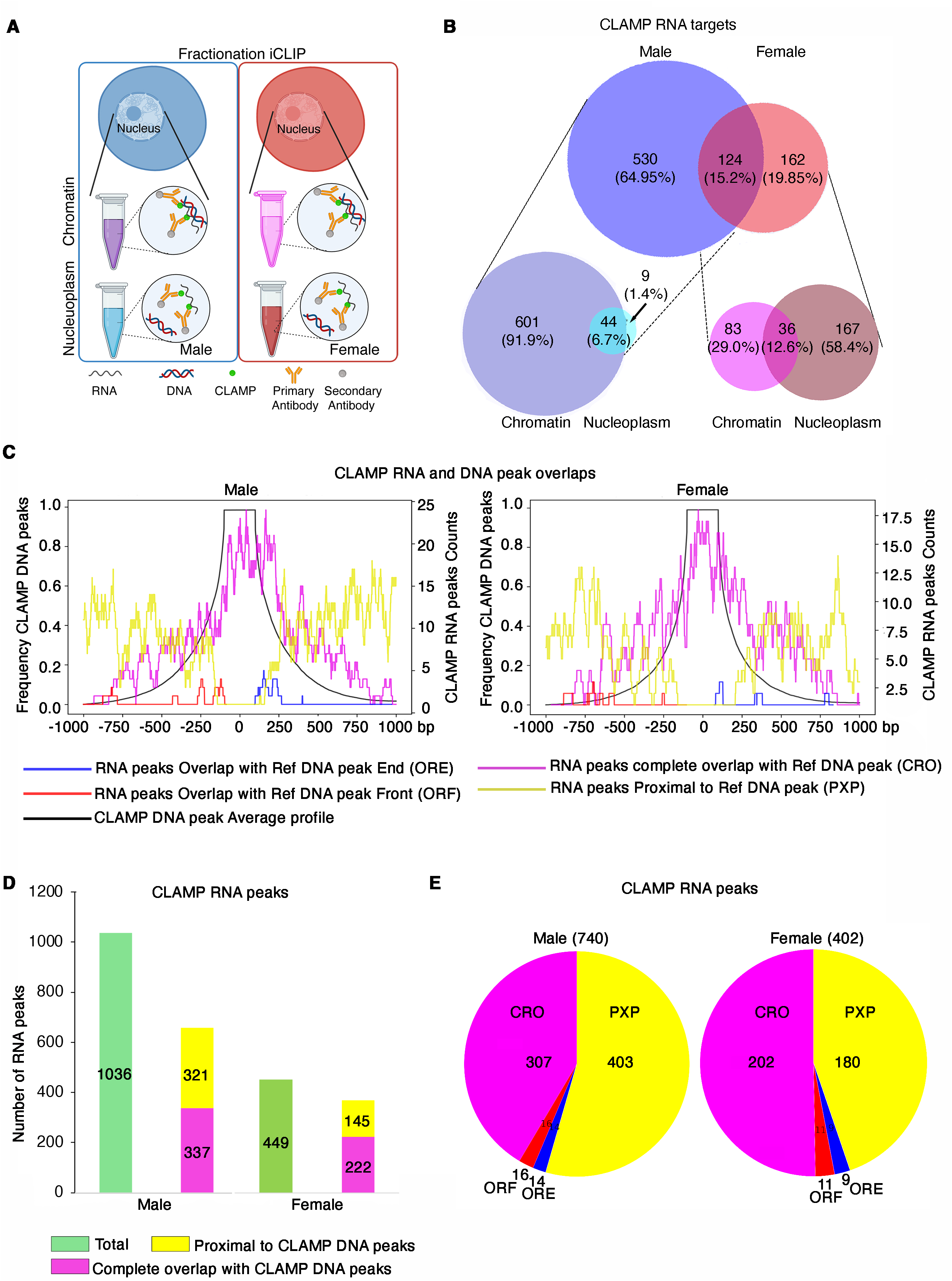
CLAMP is a dual DNA/RNA-binding protein with sex-specific RNA targets. **A.** Schematic showing the design of the fractionation iCLIP experiment to determine CLAMP RNA targets in different nuclear fractions of male and female *Drosophila* cell lines. **B**. Venn diagrams show the distribution of CLAMP RNA targets between male (S2) and female (Kc) cell types and between chromatin (ChF) and nucleoplasm (NF) fractions (**four replicates** for each category performed except **three replicates** for the Kc nucleoplasm fraction). The number within each circle/overlap region denotes the number of RNA targets/genes, and the percentage of RNA targets/gene is shown within the parentheses in each category. **C.** Frequency distribution of CLAMP RNA binding peaks represented as peak counts (right side secondary Y-axis) from iCLIP data (**four replicates** male and female chromatin fraction) plotted over a region (±1kb across the middle of the DNA peak) spanning CLAMP DNA binding peaks (black line, CUT&RUN data, **three replicates** for males, **two replicates** for females). Complete overlaps (CRO) are denoted by magenta, proximal (within ±1kb across the DNA peak),non-overlaps (PXP) in yellow, partial overlaps in red (near the starting boundary of DNA peaks, ORF), and blue (near the ending boundary of DNA peaks, ORE). **D-E.** Bar plots (**D**) and pie-chart (**E**) show the distribution of the number of CLAMP-RNA peaks that overlap with CLAMP-DNA peaks or are within a ±1kb region of CLAMP-DNA peaks (proximal peaks, PXP). Overlapping peaks are sub-categorized into complete RNA peaks overlapping with DNA peaks (CRO), overlapping RNA peaks with DNA peak front (5’ end, ORF), and overlapping RNA peaks with DNA peak 3’ end (ORE).

We found that most CLAMP interactions with RNA occur on chromatin (91.9% in males and 58.4% in females), with unique sex-specific targets, and that more RNA transcripts bind to CLAMP in males (N=654) than in females (N=286) (Fig 1B, Table 1a). Interestingly, CLAMP also directly interacts with spliceosomal RNAs sex-specifically (Table 1a). In the male chromatin fraction, CLAMP interacts with the catalytic step 2 spliceosome consisting of U2, U5, and U6 snRNAs (FDR:1.7E-3). In contrast, the female chromatin fraction is enriched for transcripts that encode proteins that bind to the U1-U2 snRNAs (FDR:1.1E-2), suggesting that CLAMP may regulate splicing differently in males and females by interacting with different spliceosomal RNAs.

Next, we asked how the CLAMP interaction with DNA correlates with its interaction with RNA on chromatin. Therefore, we used Bindcompare, a tool we developed for comparing protein binding to nucleic acids18, to plot the frequency of CLAMP RNA binding peaks on chromatin within a region ±1 kb of the closest CLAMP DNA binding peak. Using Bindcompare, we defined the following categories of binding sites: a) complete overlap (CRO) of RNA peaks with DNA peaks; b) partial overlap of RNA peak with the ends of a DNA peak (ORF or ORE); and c) RNA peaks nearby (±1 kb) DNA peaks (PXP) (Fig 1C, Fig S1a-c, Table 1b).

We found that most of the CLAMP RNA peaks identified (71.4% in males and 89.5% in females) are either proximal to (∼1 kb) or overlapping with CLAMP-DNA peaks (Fig 1D). It is important to note that the number of CLAMP-RNA peaks identified (Fig 1D) is greater than the number of RNA targets (Fig. 1B) since many targets have multiple CLAMP-binding sites (Table 1b). Notably, almost 50% of these proximal or overlapping peaks were actually within 250 bp of the middle of the nearest CLAMP DNA peak in both male and female cells, indicated by enrichment of CLAMP-RNA peaks near CLAMP-DNA peaks (Fig. 1C, E). These data suggest that CLAMP links RNA to DNA during co-transcriptional splicing at a subset of its target genes (Table 1c) that we have identified in early embryos and cell lines ^15^. Notably, only a subset of CLAMP-dependent spliced genes in the embryo and cell lines are direct CLAMP-RNA targets, revealing the context-specificity of this function. These observations reveal that CLAMP directly contacts DNA and RNA at a subset of its target genes, where it regulates splicing, indicating a direct, context-specific function in co-transcriptional splicing.

### The CLAMP prion-like domain (PrLD) is essential for CLAMP-RNA interaction

After determining that CLAMP binds RNA, we examined its domain structure to identify domains that may be responsible for RNA binding. CLAMP lacks a canonical **R**NA **r**ecognition **m**otif^18^. However, CLAMP contains a prion-like domain (PrLD), a subclass of intrinsically disordered domains (IDR) enriched in polar residues similar to those found in yeast prion proteins (Fig 1A). Many RBPs and TFs contain PrLDs^19,20^ that can promote RNA-binding and drive the formation of phase-separated biomolecular condensates^21^. Some PrLDs in TFs promote co-assembly with RBPs to regulate transcription^22–24^. However, the mechanisms by which the PrLD regions within TFs regulate alternative splicing remained unknown. Therefore, we expressed and purified full-length CLAMP protein (CLAMP^WT^-FL) and a CLAMP protein lacking the PrLD domain (CLAMP^delPrLD^-FL) (Fig 2A). We also generated the CLAMP 1-300 aa fragment, which removes the mapped DNA binding domain but includes the PrLD domain (154-300 aa). Next, we demonstrated the disordered nature of this domain (Fig 2B) using NMR spectroscopy. We also generated an NMR spectrum of a mixture of total yeast RNA extract and isolated CLAMP PrLD and found that CLAMP PrLD directly interacts with RNA (Fig 1C). Overall, we showed that the CLAMP 1-300 aa region is sufficient for direct interaction with RNA.

**Figure 2.**
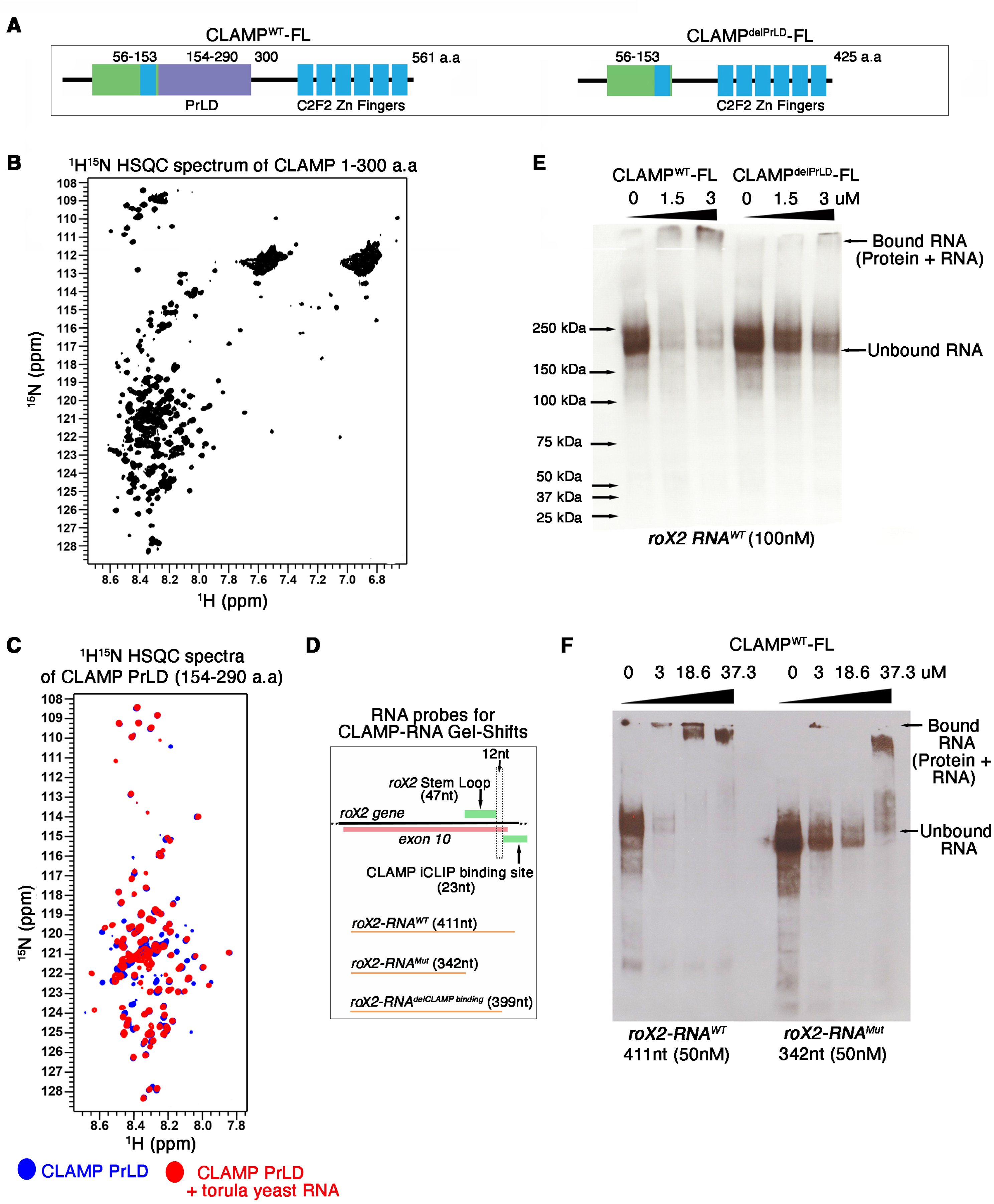
The PrLD domain of CLAMP is important for Clamp-RNA interaction. **A.** Schematic of full-length (FL) CLAMP depicting the location of the PrLD (residues 154-290 amino acids) and other essential features in the presence (left) and absence (right) of the PrLD **B.** The ^1^H^15^N HSQC spectrum of 1-300aa CLAMP at 50 µM demonstrates the largely disordered nature of the protein that lacks the DNA-binding domain. Dispersed resonances in the region from 7.9 – 7.0 ppm in the ^1^H and 115-122 ppm in the ^15^N likely arise from the folded N-terminal zinc finger **C.** ^1^H^15^N HSQC spectra of CLAMP PrLD at 20 μM with (red) and without (blue) torula yeast RNA, showing changes consistent with binding. **D.** Schematic shows the length of *roX2* RNA probes used to study the role of RNA sequence/structure in CLAMP-RNA interaction with RNA gel shift assays (RNA-EMSA). **E.** RNA-EMSA showing the binding of increasing amounts of MBP fusion CLAMP-Full length and CLAMP PrLD domain deleted protein to *roX2* RNA biotinylated probes (100 nM). Concentrations (µM) of CLAMP protein increase from left to right. **F.** RNA-EMSA showing difference in binding of full-length (FL) CLAMP (fused to MBP) to *roX2-RNA* probe with (411nt) and without (342nt) stem-loop, and CLAMP-roX2 RNA binding region identified from iCLIP. Concentrations (µM) of CLAMP FL increase from left to right. Concentrations (µM) of CLAMP FL and CLAMPdel PrLD increase from left to right.

Next, we further defined the role of the PrLD domain in promoting CLAMP-RNA interactions by performing RNA-Protein EMSA assays using specific RNA probes targeting a well-studied candidate RNA identified as a CLAMP interactor by iCLIP (Fig 2D). CLAMP is associated with the Male Sex-lethal Complex (MSL), which regulates dosage compensation in males and contains the *roX* (RNA on the X) long non-coding RNAs^13,25^. However, it was not known whether CLAMP directly interacts with the *roX* RNAs. Because the *roX* RNAs have a defined structure and function^26^, we used *roX2,* identified by iCLIP as an *in vivo* male-specific CLAMP interactor (Table 1a), to determine whether CLAMP directly interacts with RNA *in vitro*. Therefore, we used a full-length *roX2* probe (411 nt) for RNA-Protein EMSA gel shift assays (Fig 2D) with full-length CLAMP^WT^ and CLAMP^delPrLD^ proteins that were expressed and purified from *E. coli*. At the same protein and RNA concentrations, the CLAMP^delPrLD^ protein binds to RNA less efficiently than CLAMP^WT^ (Fig 2E). Therefore, the CLAMP PrLD region promotes the binding of CLAMP to RNA.

Next, we wanted to test how RNA sequence and known secondary structure regulate the CLAMP-RNA interaction. In our iCLIP experiments, we identified the CLAMP-binding region on the *roX2* RNA (23nt, Fig 2D). This region is just 12 nt downstream of a functionally important stem-loop structure (Fig 2D), which is essential for *roX2* interaction with RNA helicase A, a component of both the MSL complex and the spliceosome^26,27^. Therefore, we hypothesized that the *roX2* stem loop, known to be functionally important, promotes the interaction between roX2 RNA and CLAMP. Hence, we used two additional *roX2* RNA probes, which either completely delete both the stem-loop and the CLAMP binding region (*roX2*-RNA^mut^) or delete only the CLAMP binding region identified by iCLIP (*roX2*-RNA^delCLAMP^ ^binding^) (Fig 2D) for CLAMP-RNA Gel-shift assays.

We found that: 1) the *roX2* probe (342nt) lacking both the stem-loop and the iCLIP CLAMP binding region (69nt) reduced CLAMP binding to the *roX2* RNA (Fig 1F); 2) In contrast, the *roX2* RNA^delCLAMP^ ^binding^ probe (399nt) lacking only the iCLIP CLAMP binding region (25nt) but retaining the stem-loop, did not qualitatively affect CLAMP binding to the *roX2* RNA *in vitro* (Fig S2). Therefore, the presence of the *roX2* RNA stem-loop and the CLAMP PrLD increases the ability of CLAMP to directly interact with RNA.

### The CLAMP PrLD domain is important for CLAMP-dependent sex-specific splicing

Next, we tested whether regions of CLAMP that interact with RNA *in vitro* function *in vivo* to regulate alternative splicing. We defined the role of the CLAMP PrLD by generating complete (CLAMP Δ154-290 aa) and partial (CLAMP Δ160-290 aa) PrLD deleted mutant fly lines at the endogenous CLAMP locus using CRISPR/Cas9 genomic mutagenesis and homologous repair^39^ (Fig 3A). Complete deletion of the PrLD (*clamp^delPrLD^*) results in embryonic lethality. In contrast, homozygous partial deletion (*clamp^delPrLD+6Q^*) mutants that retain a stretch of 6 glutamines survive to adulthood, suggesting that these glutamines (aa154-160) are essential for viability (Fig 3A). Therefore, we assayed sex-specific alternative splicing in trans-heterozygous mutants (*clamp*^delPrLD^/*clamp*^delPrLD+6Q^) to ensure viability until it is possible to define the sex of animals at the larval stage (Fig 3A, B). Analysis of RNA sequencing data using the time2splice pipeline^14^ from trans-heterozygous mutants (*clamp^delPrLD^/clamp^delPrLD+6Q^*) identifies CLAMP PrLD-dependent female- and male-specific splicing (Fig 3B, Table 2a-c). More than 70% of the CLAMP PrLD domain-dependent alternative splicing was sex-specific, with ∼40-50% identified as new cryptic splicing events that do not occur in wild-type larvae (Fig 3B). Therefore, the CLAMP PrLD is essential for viability and impacts sex-specific alternative splicing.

**Figure 3.**
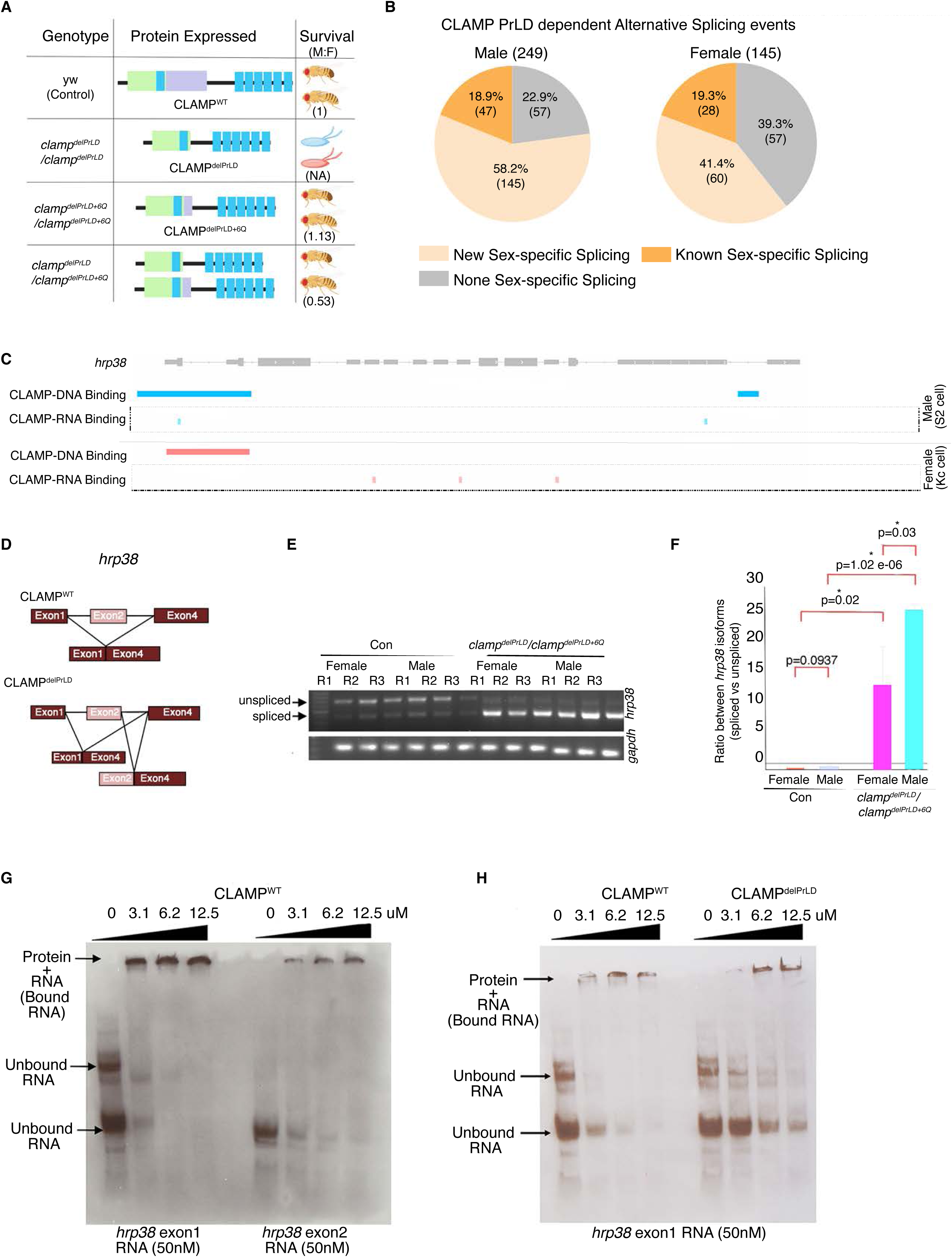
The CLAMP PrLD domain regulates sex-specific splicing in *Drosophila* third instar larvae. **A.** Region of CLAMP-Full length protein was deleted to create fly mutants expressing full (*clamp^delPrLD^*) and partial (*clamp^delPrLD+6Q^*) PrLD deleted CLAMP protein using CRISPR-Cas genome editing. The schematic shows the corresponding stages until they survive, and, within parentheses, the male-to-female ratio in adults of those that survive. **B.** Pie chart showing the proportion of CLAMP PrLD domain-dependent alternative splicing events that are sex-specific in male and female third instar larvae. The number of splicing events is noted within parentheses. **C.** CLAMP-DNA binding (CUT&RUN data) and CLAMP-RNA binding in chromatin fraction (iCLIP data) peaks visualized in the IGV genome browser at the *hrp38* gene location in female (Kc cells, red) and male (S2 cells, blue). **D.** Schematic of *hrp38* transcript showing CLAMP binding at exon1 and alternative splicing of intron AF:r6:3R:28600568:28600718:+ in *clamp*^delPrLD^/*clamp*^delPrLD+6Q^ mutant male. **E-F.** The agarose electrophoretic gel image is shown in **E,** and the corresponding bar plot in **F** shows the change in levels of intron AF:r6:3R:28600568:28600718:+ spliced isoforms resulting from alternative splicing events in male and female L3 larvae under control *clamp^WT^*(green) and *clamp*^delPrLD^/*clamp*^delPrLD+6Q^ (orange) conditions. The isoform transcript levels are normalized by the levels of the *gapdh* housekeeping gene transcript. p-values (paired Student’s t-test) for groups showing significant differences are noted at the top of the line connecting the compared groups (**three replicates** for each category). **G-H.** RNA EMSA mobility shift showing a difference in the efficiency of CLAMP full-length protein-*hrp38* exon1 and exon2 binding (**G**) and that of CLAMP-PrLD domain deleted and CLAMP full-length protein with *hrp38* exon1(**H**). CLAMP^delPrLD^ and CLAMP^WT^-Full length MBP fusions were used at increasing concentrations of 0, 3.1, 6.2, and 12.5 µM with 50 nM biotinylated RNA probes.

To determine whether the CLAMP PrLD regulates sex-specific alternative splicing of RNAs that are directly bound by CLAMP (Table 2a, b), we compared the RNAs bound to CLAMP *in vivo,* identified by iCLIP (Table 1a), with CLAMP PrLD-dependent sex-specifically spliced genes (Table 2a, b). Due to the large number of samples required, performing iCLIP in mutant L3 larvae was not possible; therefore, we used cell line iCLIP targets. We determined that 14 CLAMP PrLD-dependent sex-specifically spliced genes are direct CLAMP RNA interactors, identifying a smaller subset of key genes for further analysis (Table 3). Moreover, most of these 14 direct targets are themselves regulators of alternative splicing (Table 3). Therefore, it is possible that CLAMP, which is heavily maternally deposited^28^ and regulates splicing of 60% of all sex-specific spliced isoforms^15^, functions as an essential upstream splicing regulator by directly regulating the splicing of genes encoding splicing factors, which then regulates alternative splicing of additional target genes.

As an example of how CLAMP regulates a specific target gene involved in alternative splicing, we investigated its function relative to Hrp38, a functional orthologue of hnRNPA2B1^29–31^. Hrp38 is a strong candidate for further study because we identified it as a physical and functional interactor at the DNA, RNA, and protein levels^10^ as described below. The *hrp38* transcript is one of the 14 CLAMP PrLD-dependent spliced genes that are direct CLAMP RNA targets (Table 2a, 3). Integrating iCLIP with alternative splicing analysis suggested that in males, CLAMP binds to exon1 of the *hrp38* transcript on chromatin, in which the *hrp38* transcript undergoes splicing to remove exon2 (Fig 3C, D). Moreover, CLAMP binds both DNA and RNA at the *hrp38* gene locus (Fig 3C). As described above, CLAMP-RNA gel shift assays and NMR studies (Fig 2) indicate thatthe CLAMP PrLD domain is important for CLAMP-RNA binding. Therefore, we hypothesize that in the CLAMP *PrLD* mutant males, the mutant CLAMP protein will not bind at exon1 of the *hrp38* transcript with the same efficiency as the wildtype CLAMP (Fig 3D), which will affect splicing between exon 1 and 4, resulting in a new spliced *hrp38* RNA variant.

To test the hypothesis, we first asked whether CLAMP regulates alternative splicing at the *hrp38* locus. We found that in *clamp*^delPrLD^/*clamp*^delPrLD+6Q^ mutants, splicing of intron AF:r6:3R:28600568:28600718:+ between exon2 and exon4 (Fig 3D) is regulated by CLAMP as predicted by genome-wide splicing analysis (Table 2a). In contrast, in *clamp^delPrLD^/clamp^delPrLD+6Q^*mutants, the *hrp38* transcript retains exon2 (Fig 3E, F). Next, we validated that the alternative splicing of the *hrp38* gene is sex-specifically regulated by the CLAMP PrLD domain *in vivo* during development at the third instar larval stage in males and females. In female and male mutants, splicing of intron AF:r6:3R:28600568:28600718:+ occurs. However, in males, splicing was more efficient than in females (Fig 3F). Our *in vivo* splicing analysis thus demonstrates that the CLAMP PrLD domain is important for alternative splicing of *hrp38* during male and female development.

Since our iCLIP data show that CLAMP binds to exon1 of *hrp38* RNA, we compared CLAMP^WT^ binding to *hrp38* exon1 and exon2 probes using RNA EMSAs and found that at the same protein and RNA concentrations, CLAMP binds strongly to *hrp38* exon1 RNA with no unbound RNA remaining (Fig 3G). Furthermore, the PrLD domain of CLAMP is important for CLAMP-*hrp38* RNA binding (Fig 3H). Interestingly, CLAMP^WT^ still binds to *hrp38* exon2 *in vitro* even though it does not bind *in vivo,* suggesting that other RBPs modulate the specificity of CLAMP-RNA interactions *in vivo*. Therefore, our *in vivo* and *in vitro* approaches demonstrate that CLAMP binds to both DNA and RNA at the *hrp38* locus differentially in males and females and define how CLAMP directly regulates the sex-specific splicing of the *hrp38* transcript.

Previously, we have demonstrated using proteomics that CLAMP sex-specifically interacts with the Hrp38 protein^10^, a component of highly mobile nuclear splicing speckles that regulate alternative splicing^29,32–34^. Therefore, due to the functional interaction between CLAMP and Hrp38 at both the DNA, RNA, and protein levels, we determined how CLAMP regulates the sex-specific dynamics of Hrp38, one of CLAMP’s essential target genes and interactors, to define how CLAMP regulates sex-specific splicing beyond directly interacting with the RNA of target genes.

### CLAMP regulates the nuclear dynamics of the alternative splicing regulator hnRNPA2/B1 orthologue Hrp38

PrLD domains can drive the phase separation of proteins^21^ and promote the formation of subnuclear bio-condensates involved in transcription^22,35^ and splicing^21,36^. Recent advances show that biophysical properties of biomolecular condensates regulate splicing function^37^. Therefore, we performed biochemical phase-separation assays to determine how the CLAMP PrLD regulates CLAMP phase separation (Fig 4A). We found that the CLAMP PrLD contributes to phase separation of the N-terminal half of CLAMP (residues 1-300). Many RBPs involved in splicing form splicing condensates, and TFs form transcription condensates, but the mechanisms by which transcription and splicing condensates interact to promote the fidelity of co-transcriptional splicing remain very poorly understood^38^. Although the PrLDs of TFs can promote co-assembly of biomolecular condensates with RBPs6,22,40, current work does not yet explain how TF-RBP interactions regulate the dynamics of splicing condensates, which are linked to their function^37,39^.

**Figure 4.**
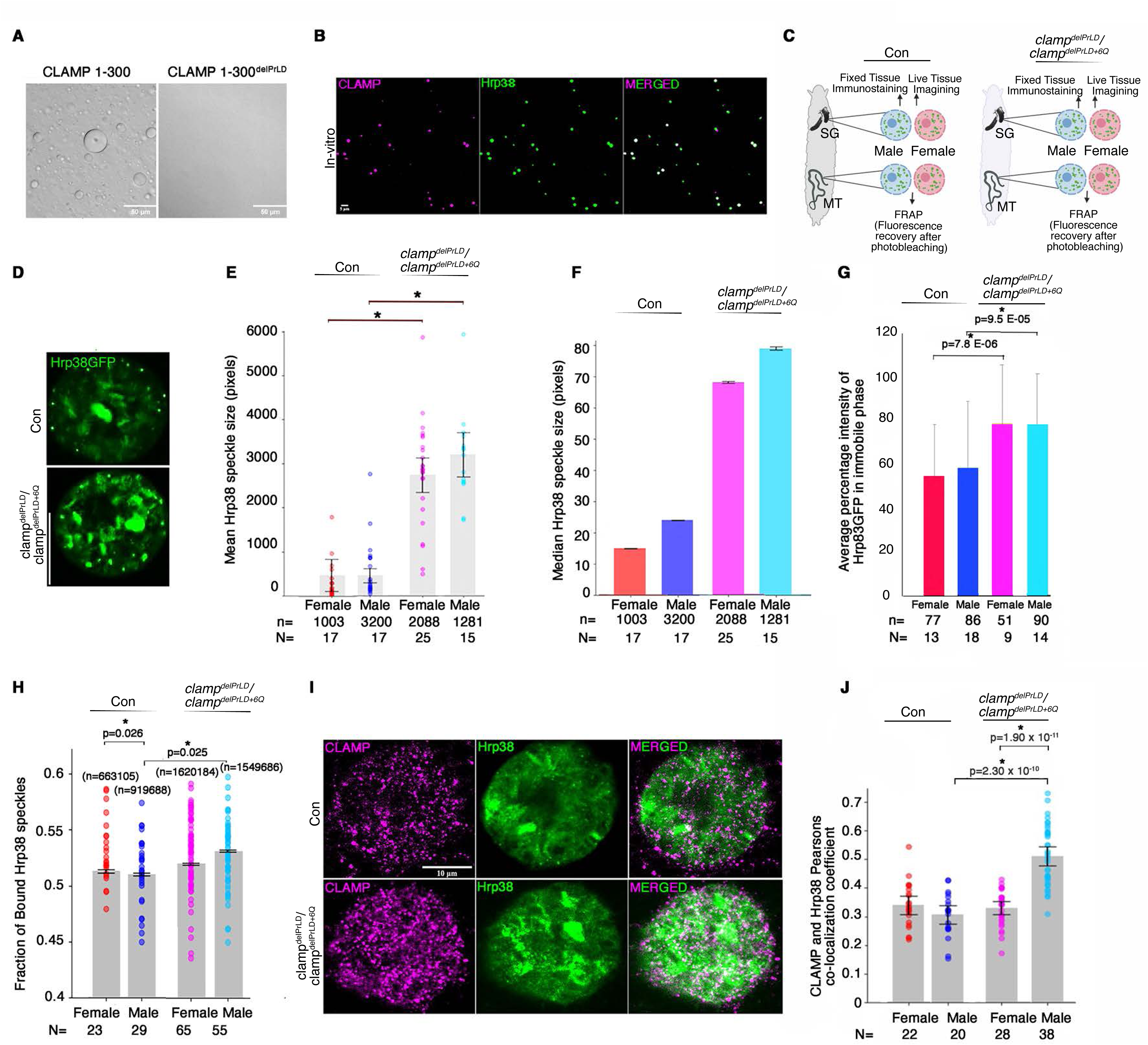
The CLAMP PrLD domain regulates the dynamics of hnRNPA2 homolog Hrp38 biomolecular condensates. **A**. DIC micrograph of CLAMP 1-300 with or without PrLD domain (100 µM) at 150 mM salt concentration with 5% PEG. **B**. Single plane confocal images showing the co-phase separation of Alexa Fluor 488 maleimide dye-labeled Hrp38 with a single cysteine added for labeling (green) and Alexa Fluor 555 maleimide dye-labeled 1-300 CLAMP (pseudo color, magenta) under phase separating conditions for Hrp38. **C**. Schematic showing experimental design setup for studying Hrp38 nuclear condensates in vivo using fixed and live tissue imaging techniques. **D-F.** Confocal image of a single plane of third instar larval salivary gland nuclei expressing Hrp83-GFP in *clamp^WT^*(green) and *clamp*^delPrLD^/*clamp*^delPrLD+6Q^ conditions in **D,** with bar plots showing mean (**E**) and median (**F**) Hrp38 speckle/condensate size in control and *clamp*^delPrLD^/*clamp*^delPrLD+6Q^ male and females. p-values (paired Student’s t-test) for groups showing significant differences in mean speckle size are noted at the top of the line connecting the compared groups. **G**. Bar plots showing the percentage of Hrp38-GFP in the immobile phase (FRAP analysis) in male and female L3 larvae under control *clamp^WT^* and *clamp*^delPrLD^/*clamp*^delPrLD+6Q^ conditions. p-values (paired Student’s t-test) for groups showing significant differences marked by an asterisk are noted at the top of the line connecting the compared groups—**n=**Number of Malpighian tubule principal cell nuclei, **N=** Number of individuals. **H**. Mean bound fraction of Hrp38GFP speckles (Number of Hrp38GFP speckles bound, i.e., immobile track lengths/the total number of Hrp38GFP speckles with all types of track lengths over a fixed time-period) from live imaging of salivary gland nuclei expressing Hrp38-GFP in male and female *clamp^WT^*and *clamp*^delPrLD^/*clamp*^delPrLD+6Q^ individuals. Each dot **N** represents each individual (movie), and ‘**n**’ denotes the number of Hrp38 speckles identified in each category. p-values (paired Student’s t-test) for groups showing significant differences in the mean fraction of bound Hrp38 speckles are noted at the top of the line connecting the compared groups. **I-J.** Confocal single plane images of immune-stained third instar larval salivary gland nuclei showing in vivo partial co-localization of CLAMP (magenta) and Hrp38 (green) speckles in control and mutant (*clamp*^delPrLD^/*clamp*^delPrLD+6Q^) males in **I** with corresponding bar plots in **J** showing the Pearson’s colocalization coefficient between CLAMP (magenta) and Hrp38 (green) in both males and females. p-values (paired Student’s t-test) for groups showing significant differences are noted at the top of the line connecting the compared groups.

The hnRNPA1/B2 orthologue Hrp38 is a strong candidate to function with CLAMP to promote alternative splicing because it is associated with CLAMP in three ways: 1) CLAMP interacts with the *hrp38* RNA transcript in our iCLIP experiments (Table 1a); 2) CLAMP regulates alternative splicing of the *hrp38* transcript (Table 2, Fig 3), and 3) CLAMP interacts with the Hrp38 protein sex-specifically^10^. Hrp38 forms dynamic nuclear splicing condensates or speckles^29,32–34^. Therefore, we hypothesized that CLAMP PrLD-mediated phase separation affects the dynamics of Hrp38 condensates. We tested our hypothesis by combining *in vitro* and *in vivo* approaches to measure condensate formation, stability, and dynamics.

In *vitro*, CLAMP 1-300^WT^ and Hrp38 co-phase-separate into liquid-like condensates (Fig 4B). To complement *in vitro* studies, we performed *in vivo* analysis of Hrp38 nuclear speckles in fixed and live conditions in *Drosophila* third instar larval tissue (Fig 4C). Salivary gland (SG) and malpighian tubule (MT) tissue nuclei were used since they have large polyploid or polytene nuclei, making it easier to visualize and track nuclear speckles. Prior studies^40–43^ suggest that larger aggregates of hnRNPs are more immobile and have decreased function in regulating alternative splicing. Therefore, we performed the following experiments in males and females to test our hypothesis that the CLAMP PrLD domain regulates the dynamics of Hrp38 to drive changes in sex-specific splicing: 1) We analyzed fixed SG nuclei expressing Hrp38-GFP samples to determine the effect of the CLAMP PrLD domain on Hrp38 nuclear speckle size (Fig 4C-F); 2) We performed live imaging to define how the CLAMP PrLD domain regulates the mobility of Hrp38-GFP nuclear speckles (Fig 4C, G); 3) We performed FRAP analysis in MT nuclei to measure the mobility of the Hrp38 protein in the nucleus by measuring the fraction of Hrp38 that remains bound (Fig 4C, H). We also analyzed the co-localization of CLAMP and Hrp38 *in vivo* (Fig. 4I-J) to determine how frequently they interact, and compared these findings with our in vitro interaction results (Fig. 4B).

Compared with wild-type controls, in *clamp^delPrLD^* mutants, we observe larger Hrp38 speckles (Fig. 4D-F) with lower mobility (Figs. 4G, H). The Hrp38 protein forms nuclear speckles (Fig 4D), which are highly dynamic (Movie 1); however, in clampdelPrLD/clampdelPrLD+6Q mutants, Hrp38 forms larger nuclear aggregates in both females and males compared to controls (Fig 4E), with median speckle size in males slightly higher than in females (Figure 4F). Hrp38 nuclear size negatively correlated with speckle dynamics ( Fig 4G, H, Movie 2). FRAP analysis shows that in *clamp*^delPrLD^/*clamp*^delPrLD+6Q^ mutant male and female individuals, Hrp38 remains more immobile compared with matched wild-type controls (Fig 4G, Movie 3,4). Therefore, the presence of the CLAMP PrLD domain reduces the size and increases the dynamics of the Hrp38 speckles.

Next, we analyzed the trajectories of moving Hrp38 speckles over time with live image analysis (Materials and Methods) to determine whether CLAMP regulates their mobility. More restricted tracks denote more immobile speckles, while more dispersed tracks represent more mobile speckles^44^. Interestingly, we found that in wild-type controls, female Hrp38 speckles are less mobile than male Hrp38 speckles. Without the CLAMP PrLD domain, Hrp38 speckles in males become less mobile (Fig 4H), therefore becoming more similar to those in females. Therefore, the CLAMP PrLD regulates the dynamics of Hrp38 speckles sex-specifically in live tissues, promoting their enhanced mobility in males. We identified partial co-localization between CLAMP and Hrp38, which is significantly increased in *clamp^delPrLD^*mutant males (Fig. 4I- J). Therefore, the CLAMP PrLD domain in males also prevents colocalization with Hrp38, supporting its differential role in influencing Hrp38 dynamics in males and females.

Thus, the presence of the CLAMP PrLD reduces the size of Hrp38 splicing condensates, which has been shown to increase their ability to mediate alternative splicing with high fidelity^45,46^ Therefore, our data support a model in which the CLAMP PrLD modulates Hrp38 condensates’ ability to regulate alternative splicing. Furthermore, the CLAMP PrLD promotes the formation of CLAMP liquid droplets (Fig 4A), which can co-localize with Hrp38 splicing condensates (Fig. 4B). Deletion of CLAMP PrLD inhibits CLAMP phase separation *in vitro* (Fig 4A) and sex-specifically regulates the dynamics of Hrp38 speckles *in vivo* (Fig. 4G- H). Overall, our data suggest that the CLAMP PrLD sex-specifically alters the dynamic properties of Hrp38, preventing it from forming large aggregates, which have been demonstrated to be non-functional^45,46^.

Our data show that the PrLD domain of CLAMP is important for its phase-separation properties (Fig 4A) and regulates the dynamics of the Hrp38 condensate under cellular conditions (Fig. 4D-H). In vitro data also support the idea that CLAMP and Hrp38 can coexist as phase-separating condensates (Fig 4B). Interestingly, in females, where Hrp38 protein associates with CLAMP^10^, Hrp38 protein is less dynamic and more in bound form compared to males (Fig 4H). The PrLD domain of CLAMP plays an important role in the male-specific dynamic physical properties of Hrp38. In *clamp*^delPrLD^/*clamp*^delPrLD+6Q^ mutants, there is an increase in bound Hrp38 and slower Hrp38 dynamics in males (Fig 4H). Since this is associated with enhanced CLAMP and Hrp38 protein co-localization *in vivo* in *clamp*^delPrLD^/*clamp*^delPrLD+6Q^ males, we propose a hypothesis that CLAMP and Hrp38 association in females and a lack of the same in males, mediated by the PrLD domain of CLAMP, results in a difference in Hrp38 dynamics between the two sexes (Fig 4H). The effect this difference has on Hrp38 function and its regulation needs further investigation. Currently, our working model highlights that an Intrinsically disordered Domain (IDR), such as PrLD in CLAMP (TF), changes biophysical properties of partner proteins (RBPs), drives sex-specific protein-protein (Fig. 4I-J) and protein-RNA association (Fig. 2E, 3H), resulting in sex-specific splicing changes (Fig. 3B, E-F).

## Discussion

Very little is understood about how alternative splicing is targeted to the right place at the right time within a highly compacted genome to prevent the formation of cryptic splicing events that contribute to disease^1^. We hypothesized that transcription factors that interact with RNA-binding proteins would be strong candidates for targeting alternative splicing to the correct genomic locations in a developmental context. To test our hypothesis, we examined the *Drosophila* CLAMP TF as a model to reveal new mechanisms by which the hundreds of TFs interacting with RNA and RNA-binding proteins regulate splicing, because it regulates a well-defined alternative splicing process, sex-specific splicing^15^, and we can integrate genetic, genomic, live imaging, biochemical, and structural approaches.

CLAMP is a pioneer TF that interacts with the male-specific MSL complex to regulate *Drosophila* dosage compensation. It is essential for MSL targeting to GA-rich chromatin entry sites (CES) on chromatin, thus helping to target the MSL complex to the X chromosome. Interestingly, CES evolved from intronic polypyrimidine tracts, suggesting a further link with splicing^47^. We found that, mechanistically similar to its role in dosage compensation, where it recruits a ribonucleoprotein complex containing the *roX* non-coding RNAs, CLAMP directly links RNA to DNA at a subset of target genes where it regulates sex-specific splicing.

Because not all CLAMP-dependent sex-specific spliced genes are both direct CLAMP-RNA and CLAMP-DNA targets, CLAMP regulates alternative splicing through multiple mechanisms that may be regulated by the presence of MSL complex in males but not females. Interestingly, CLAMP regulates the splicing of many RBPs that are components of alternative splicing complexes, including *hrp38.* The alternatively spliced forms of these proteins can result in protein isoforms with different functions. CLAMP-regulated alternate first (AF) alternative splicing of *hrp38* transcript gives rise to two different protein isoforms (Fig 3). How these two Hrp38 protein isoforms differ in their physical and functional properties and how they regulate sex-specific splicing is an important direction for future work.

We define CLAMP-regulated RBP protein dynamics as an alternative mechanism by which CLAMP regulates splicing. We show that the CLAMP PrLD domain is essential for determining the size of Hrp38 protein condensates, which in turn regulates their function. We hypothesize that CLAMP affects Hrp38 splicing speckles by regulating the biophysical phase separation properties of Hrp38 condensates. Aggregates of human Hrp38 homologue (HnRNPA2B1) results in disfunction and disease phenotype, indicating the right size of the phase separated condensates is essential for its function^45,46^. We show that CLAMP PrLD domain is essential for the Hrp38 condensate size, its loss resulting in aggregation (Fig. 4D-F), loss of phase separation properties and dynamics (Fig. 4G-H) of Hrp38 protein, and would ultimately affect its function. Our data support a model in which the CLAMP PrLD domain differentially regulates CLAMP-Hrp38 interactions to regulate splicing condensate properties in males and females. We further propose that the interaction of CLAMP with the MSL complex in males but not females promotes sex differences in Hrp38 function. The following evidence supports this model: 1) Hrp38 mobility is different between males and females (Fig 4J) as is the CLAMP-Hrp38 protein interaction which we identified as female-enriched from proteomic analysis^10^; 2) Hrp38 splicing condensate mobility is more dysregulated males than in females in *clamp PrLD* mutants (Fig 4J); 3) The colocalization between CLAMP and Hrp38 is decreased in males versus females, potentially due to competition from the CLAMP-MSL interaction (Fig. 4I- J). Overall, our data support a model in which CLAMP sex-specifically regulates the dynamics of the hnRNP orthologue Hrp38.

Thus, one of the determinants of the sex-specificity of the CLAMP-dependent alternative splicing is sex-dependent variability in Hrp38 condensate dynamics. CLAMP interacts with the Hrp38 protein differentially between sexes^10^. Also, the PrLD domain of CLAMP regulates Hrp38 condensates more strongly in males, predicting that the CLAMP PrLD domain is important for these male-biased interactions. This might be due to CLAMP interacting with the MSL complex in males, thereby inhibiting its interaction with other proteins, such as Hrp38. However, the absence of this complex would make such associations possible. The biomolecular interactions by which the PrLD domain of CLAMP regulates Hrp38, determining CLAMP-Hrp38 interactions needs further study.

Another determinant of the sex-specificity of CLAMP-dependent alternative splicing is that CLAMP has different RNA targets in males and females, with the PrLD domain again playing an important role. We hypothesize that sex-specific CLAMP-RNA targets result from CLAMP having different RBP interactors in males and females, especially closer to chromatin^10^. his might result from differential interactions between CLAMP and different ribonucleoprotein complexes such as the MSL complex (MSL proteins and *roX* RNA) or the spliceosomal complex (RBPs and snRNAs) on chromatin. In the future, new techniques like RD-SPRITE^48^ that identify the distribution of different nuclear clusters of RNAs on chromatin, including splicing hubs, can be used to determine the global as well as specific local roles of CLAMP in establishing site-specific RNA splicing in male compared with female cells.

We found substantial new cryptic splicing events^1^ occurring in the absence of CLAMP (Fig 3B), which indicates that at most genomic sites, CLAMP acts as an inhibitor of cryptic alternative splicing events. Thus, overall, our data support a model where sex-specific differences in alternative splicing arise from differential TF-RNA and TF-RBP interactions (Figs. 1 and 4). We show for the first time that an intrinsically disordered domain (IDR) like the PrLD domain of CLAMP in a TF regulates alternative splicing by altering the dynamics of splicing condensates. Also, our data suggest that TFs that bind to specific DNA motifs and have RNA binding properties are strong candidates for understanding how specific alternative splicing events are regulated context-specifically to drive normal development and prevent the onset of disease states.

## Methods

### Cell culture

Kc and S2 cells were maintained at 25°C in Schneider’s media supplemented with 10% Fetal Bovine Serum and 1.4X Antibiotic-Antimycotic (Thermofisher Scientific, USA). Cells were passaged every three days to maintain an appropriate cell density.

### Fly stocks and husbandry

*Drosophila melanogaster* fly stocks were maintained at 24°C on standard corn flour sucrose media. Fly strains with complete (CLAMP Δ154-290a.a) and partial (CLAMP Δ160-290a.a) PrLD deleted mutant fly lines using CRISPR/Cas9 genomic mutagenesis and homologous repair^49^. We used the flyCRISPR Optimal Target Finder tool from the University of Wisconsin to design a CRISPR target sequence for *clamp^delPrLD^*and *clamp^delPrLD+6Q39^*. We cloned target sequence oligonucleotides (one gRNA) for *clamp^delPrLD^* (sense:5’CTTCGGGTACAACGCCAAAGCGAG3’;antisense:3’CCCATGTTGCGGTTTCGCTC -CAAA5’) and two gRNAs for *clamp^delPrLD+6Q^* (sense: 5’CTTCG-AATAGAATCGCCGCCCGCT3’;antisense:3’CTTATCTTAGCGGCGGGCGACAAA5’)and(sense:5’CTTC-GTTGTGGCTGCACAGACTGG3’;antisense:3’CAACACCGACGTGTCTGACC-CAAA5’) into the pU6-BbsI-chiRNA plasmid (Addgene no. 45946), following the protocol outlined on the flyCRISPR website. We validated the correct ligation of the *clamp* CRISPR target sequence into the pU6-BbsI-chiRNA plasmid by Sanger sequencing using universal M13 primers. For homologous repair, we used ssODN15’ACATAAGCTTTAAGTGTGACGTATGTTCAGATATGTTCCCTCATTTGGCACTTCTTAATGCTCATAGAGAGGCGAGCAGTGGGTCTGGCCATCATCCTGTGAAAAAACGAAATTCCCAGCAGATGACCAAAT3’ for *clamp^delPrLD^*and ssODN2 5’CTTCTTAATGCTCATAAGCGGATGCATACAGACGGGGAACAGCAGCAACAACAGCAACATAACGCCCAAGCTGGCGGTACAACGCCAAAGCGAGAGGCGAGCAGTGGGTCTGGCCATCATCCTGTGAAAAAA3’ for *clamp^delPrLD+6Q^*.The commercial service, BestGene Inc., microinjected the validated pU6-BbsI-chiRNA plasmid containing the *clamp* target sequence into germline-expressing Cas9 flies Bloomington stock #51324. Flies containing a single mutation were returned balanced over the *Curly of Oster* (*CyO)* second chromosome balancer. From these progenies, we identified the CRISPR/Cas9-generated mutations by PCR across the target region (forward:5′-GATATGTTCCCTCATTTGGCAC-3′, reverse:5′-CACTCCCATGCTTCACACAG-3′). We isolated two independent clamp alleles from this validation: (1) *y^1^*, *w^1118^*; *clamp^delPrLD^*/*CyO*; and (2) *y^1^*, *w^1118^*; *clamp^delPrLD+6Q^*/*CyO.* These were crossed to obtain male and female *clamp^delPrLD+6Q^*/*clamp^delPrLD^* genotypes. Crisper/Cas9 generated fly mutants and fly strain y^1^ w^1118^; P{w[+mC] =PTT-GC} Hrb98DE[ZCL0588] expressing Hrp38GFP (Bloomington #6822) were crossed to obtain *clamp^delPrLD+6Q^*/*clamp^delPrLD^*male and female expressing GFP tagged Hrp38 protein.

### CUT&RUN in cell lines

Cells were allowed to grow to confluency and harvested. An equal number of cells for each category was suspended in wash buffer and subjected to the Cut&Run assay according to Skene et al. 2018^50^ using rabbit anti-CLAMP (5µg) to immunoprecipitate CLAMP-bound DNA fragments from male (S2) and female (Kc) cell lines. Three replicates for males and females were run, but one female sample was dropped during later stages due to insufficient starting material. Rabbit IgG was used as a control for each male and female cell line sample. 1ng CUT&RUN DNA was used to generate libraries using the Kapa Hyper prep kit (Roche, USA) and SeqCapAdapter Kit A (Roche, USA). 14 PCR cycles were used to amplify the libraries. AMPure XP beads (Beckman Coulter, USA) were used for library purification, and fragment analysis was performed to check the library quality. Paired-end 2x25 bp Illumina Hi-seq sequencing was performed.

### RNA-seq in third instar larvae (L3)

Total RNA was extracted from control (y^1^ w^1118^; *hrp38GFP*) and *clamp* mutant (*yw, clamp^delPrLD+6Q^*/*clamp^delPrLD^*; *hrp38GFP*) male and female third instar larvae (3 each) using Trizol (Invitrogen, USA). Four replicates were in each category prepared. Messenger RNA was purified from total RNA using poly-T oligo-attached magnetic beads. After fragmentation, the first strand of cDNA was synthesized using random hexamer primers, followed by the second cDNA synthesis. The library was ready after end repair, A-tailing, adapter ligation, size selection, amplification, and purification, followed by paired-end RNA-sequencing in Illumina Novaseq 6000. The sequencing data were run through a SUPPA-based time2splice pipeline^15^ to identify CLAMP-dependent sex-specific splicing events. Data is to be submitted to the GEO repository.

### iCLIP

Cells were allowed to grow to confluency, and UV crosslinked using 254 nm UV light in Stratalinker 2400 on ice (Stratagene, USA). UV-treated cells were lysed to get different cellular fractions (Cytoplasmic, Nucleoplasmic, and Chromatin) according to the Fr-iCLIP (fractionation-iCLIP) protocol from Brugiolo et al 2017^51^. Chromatin and nucleoplasmic fractions were sonicated with a Branson digital sonicator at 30% amplitude for 30 seconds (10 sec on, 20 sec off) to disrupt DNA prior to IP. All three fractions were separately centrifuged at 20,000 xg for 5 min at 4℃. Fractions were tested by Western blotting using RNApolI for the Chromatin Fraction and Actin for the Cytoplasmic Fraction. Protein quantification for each fraction was done using the manufacturer’s protocol for Pierce 660 nm protein assay reagent (Thermo Scientific, USA). Each Fraction was subjected to the iCLIP protocol as described in Huppertz et al. 2014^52^ using the rabbit-CLAMP antibody to immunoprecipitate bound RNAs extracted using proteinase K and phenol: chloroform. Custom cDNA libraries were prepared according to Huppertz et al. 2014^52^ using distinct primers Rt1clip-Rt16clip for separate samples containing individual 4nt-barcode sequences that allow multiplexing of samples. cDNA libraries for each sample were amplified separately using 31 cycles of PCR, mixed later, and sequenced using standard Illumina protocols. Heyl et al. 2020^53^ methods using the Galaxy CLIP-Explorer were followed to preprocess, perform quality control, post-process, and peak calling.

### Validation of hrp38 splicing using RT-PCR assay

Total RNA was extracted from 5 third instar larvae (L3) female and male embryos expressing *clamp^delPrLD+6Q^*/*clamp^delPrLD^*and *y^1^*, *w^1118^*. Following the manufacturer’s protocol, we reverse-transcribed one microgram of total RNA using the SuperScript VILO cDNA Synthesis Kit (Life Technologies, USA). We amplified target sequences by PCR using primers designed to span alternatively spliced junctions (FP-5’AGAACGGCAACTCCAATGGC3’ and RP-5’GCCAGTCTCCTTGTCAATGA3’) and Quick load Taq 2X Master mix (#M0271L, NEB, USA) according to the manufacturer’s protocol (28 cycles). 10ul of PCR product of each replicate for each gene was loaded in separate wells in 2% agarose gels and imaged using a ChemiDoc^TM^ MP Imaging system (BioRad, USA). All replicates for each gene were loaded on the same gel. The gel images were quantified using the densitometry steps with the Fiji image analysis tool. Student’s t-tests were performed to determine significant differences between groups (two samples at a time). Three replicates for RT-PCR samples were performed.

### CLAMP Protein Expression and Purification

Vectors encoding maltose binding protein (MBP)-tagged CLAMP 1-300 and MBP-tagged CLAMP 1-300 delPrLD were produced by cloning into the pTHMT vector. Plasmids were transformed into BL21 Star (DE3) cells, and bacterial cultures were grown in a minimal M9 medium supplemented with ^15^N ammonium chloride. Cultures were grown at 37°C and 200 RPM to an optical density of 0.6-0.8 and subsequently induced with 1 mM isopropyl-β-D-1-thiogalactopyranoside (IPTG) for 4 hours at 37°C. Cell pellets were harvested by centrifugation at 6,000 RPM, resuspended in lysis buffer (20 mM Tris pH 8, 1 M NaCl, 10 mM Imidazole, 1 mM DTT, and one EDTA-free protease inhibitor tablet (Roche) per liter of culture), and lysed by sonication. The lysed cell suspension was cleared by centrifugation at 20,000 RPM for 50 min. The supernatant was filtered using a 0.2 μm syringe filter and loaded onto a HisTrap HP 5 ml column. The HisTrap column was first washed with 20 mM Tris pH 8, 1 M NaCl, 10 mM Imidazole, 1 mM DTT Buffer, and then the bound protein was eluted with 20 mM Tris pH 8, 1 M NaCl, 500 mM Imidazole, 1 mM DTT buffer. Fractions containing MBP-CLAMP 1-300 were collected and purified on a Superdex 200 (26/600) column equilibrated in 20 mM Tris pH 8, 1.0 M NaCl buffer. As above, CLAMP PrLD (154-290) was expressed and purified using HisTrap HP. Following that, CLAMP PrLD was collected from the HisTrap HP, concentrated to 1 mL, and cleaved overnight with TEV protease to separate CLAMP PrLD from the MBP in 20 mM Tris pH 8, 1 M NaCl, 10 mM Imidazole, and 1 mM DTT Buffer. The cleaved protein was separated from the HisTag by a subtraction step, in which the cleaved protein was flowed over the HisTrap HP column using the same Tris buffer previously described, and the flow-through was collected. MBP-tagged Full-Length CLAMP was grown following the same methods above. Once the Full-Length CLAMP cell lysate supernatant was loaded onto the HisTrap HP 5 ml column, the protein was eluted with a gradient from 10 to 500 mM imidazole in 20 mM MES at pH 5.5 and 361 mM CaCl_2_. Fractions containing MBP-CLAMP FL were collected and purified on a Superdex 200 (26/600) column equilibrated in pH 5.5, 20 mM MES, and 361 mM CaCl2 buffer. For each CLAMP construct, fractions containing the desired protein were verified by SDS-PAGE, concentrated using a 10 kDa centrifugation filter (Millipore), aliquoted, and flash-frozen in liquid nitrogen.

### Hrp38 protein purification and phase separation assay

Maltose-binding protein (MBP)-tagged Hrp38 was expressed in BL21 Star (DE3) *E. coli* cells. The cells were resuspended in 20 mM NaPi at pH 7.4 with 300 mM NaCl and 10 mM imidazole, and the lysate was cleared by centrifuging at 20,000 rpm for 60 mins at 4 °C. The supernatant was filtered using 0.2 µM filters and loaded on a 5 ml HisTrap HP column. The protein was eluted using an imidazole gradient of 10-300 mM over 5-column volumes. The protein fractions were pooled and loaded on a Superdex 200 (26/600) column for size exclusion. 20 mM NaPi with 300 mM NaCl at pH 7.4 was used for size exclusion chromatography and storage. Protein was flash frozen as aliquots of 1 mM. Fluorescent labeling of the CLAMP 1-300 was done using AlexaFluor 488 maleimide dye, while Hrp38 was labeled using 555 maleimide. The protein was diluted to 100 µM in 20 mM Tris buffer at pH 7.4 with 50 mM NaCl, and a 5-fold concentration (500 µM) of the dye dissolved in DMSO was added to 5% of the total volume. The reaction mixture was incubated for 1 hour, and then, unbound AlexaFluor was removed using 1 ml Zeba spin desalting columns. The proteins were concentrated to 1 mM, flash frozen, and stored. For phase separation and microscopy, Hrp38 was buffer-exchanged into 20 mM Tris buffer at pH 7.4 containing 50 mM NaCl to a final concentration of 20 µM. Stock and sample concentrations were measured using a NanoDrop spectrophotometer with extinction coefficients of 130,000 M^-1^ cm^-1^ for Hrp38 and 78270 for MBP CLAMP 1-300. 1 μM of AlexaFluor-labeled CLAMP was added to Hrp38, and TEV protease was added to cleave the MBP tag. Less than 1% fluorescent protein was used in all samples. A Nikon spinning-disc confocal microscope (Nikon Eclipse Ti2) was used for imaging at 20X magnification with 1.5X zoom. The images were processed using ImageJ.

### NMR Sample Preparation and Spectroscopy

^15^N isotopically labeled samples of CLAMP 1-300 were prepared at 50 µM in a buffer containing 20 mM MES pH 5.5, 100 mM NaCl, 1 mM DTT, and 5% D_2_O, and then moved into a 5 mm NMR tube using a glass pipette. CLAMP concentration was determined by measuring absorbance at 280 nm (and then dividing absorbance by the extinction coefficient estimated by the Expasy ProtParam). NMR spectra were recorded ona Bruker Avance 850 MHz ^1^H Larmor frequency spectrometer with HCN TCI z-gradient cryoprobes. A two-dimensional ^1^H^15^N HSQC was acquired using spectral widths of 10.5 ppm and 30.0 ppm in the direct and indirect dimensions, with 3072 and 512 total points and acquisition times of 172 ms and 99.0 ms, respectively. Samples of isotopically labeled CLAMP PrLD were prepared at 20 µM in a buffer containing 50 mM MES, pH 5, and 5% D_2_O. Because the PrLD domain of CLAMP contains no tyrosines or tryptophans, the A_280_ absorbance could not be measured to determine the protein concentration. Instead, sample concentration was estimated by measuring absorbance at 230 nm and calculating the extinction coefficient to be 300 M-1 cm-1 per peptide bond (40.5 mM-1 cm-1 for CLAMP PrLD). Torula yeast was added to one PrLD sample at a 1:1 Protein to RNA ratio by weight, and NMR spectra were acquired in 5 mm NMR tubes on a Bruker Avance 850 MHz ^1^H Larmor frequency spectrometer. Both 1H15N HSQC spectra were acquired with spectral widths of 14.0 ppm and 25.0 ppm in the direct and indirect dimensions, respectively, with 2048 and 256 total points and acquisition times of 86.0 ms and 59.4 ms, respectively. All data were processed and analyzed using NMRPipe and CCPNMR Analysis 2.5 software^54,55^.

### RNA Electrophoretic Mobility Shift Assays

*rox2*RNA probes at 100 nM and *hrp38* RNA probes at 50nM were incubated with MBP-tagged FL CLAMP^WT^ protein or MBP-tagged FL CLAMP^delPrLD^ protein in REMSA binding buffer provided with the LightShift Chemiluminescent RNA EMSA kit (Thermo Scientific, USA) at room temperature for 30 min according to the manufacturer’s protocol. Reactions were loaded onto 6% TBE retardation gels (Thermo Fisher Scientific) and run in 0.5× Tris–borate–EDTA buffer for one hour. RNA-Protein complex was transferred to the Nylon membrane using the iBlot transfer system (ThermoFisher Scientific), and the probe signal was detected using a Chemiluminescent Nucleic acid detection module (#80880, ThermoFisher Scientific).

### Immunostaining of *Drosophila* tissue

Third instar larval salivary gland tissue from control and *clamp^delPrLD+6Q^*/*clamp^delPrLD^* male and female dissected in 1XPBS (Phosphate Buffer Saline, 10X PBS, Boston Bioproducts Inc, USA) and fixed in 4% PFA in 1XPBS (Paraformaldehyde, Sigma Aldrich, USA) for 20 min. Incubated in 1% PBST (Triton-X in 1XPBS) for 10 minutes and washed twice in 0.1% PBST for 10 minutes each. They were incubated in a moist chamber with a blocking solution (3% BSA, bovine serum albumin in 0.1%PBST) for 2 hours at RT. Tissues were then incubated in rabbit anti-CLAMP (1:500, SDIX) overnight at 4 degrees Celsius in a blocking solution in a moist chamber. Washed tissues in 0.1% PBST three times for 15 minutes each. Secondary anti-rabbit Alexa fluor 568 at 1:200 dilution was used to incubate tissue for 2 hrs at RT in a blocking solution in a moist chamber and again, washed in 0.1% PBST three times for 15 minutes each, followed by DAPI staining (1ug/ml in 1XPBS) for 10 min at RT in the dark. After two washes in 0.1% PBST for 10 min each, tissues were mounted on a slide in Vectashield (Vector Laboratories Inc, USA).

### Fluorescence recovery after photobleaching (FRAP)

*In vivo* (Malphighian tubule principal cell nucleus expressing Hrp38GFP in control and *clamp^delPrLD+6Q^*/*clamp^delPrLD^*male and female third instar larvae) FRAP was performed on a Nikon Spin-disc Confocal Microscope with a 488 nm laser on a 60× objective,taking frames without delay (short time course and fast-recovering control) with 3-sec acquisition pre-bleaching, bleaching using 473 nm laser at 100%, and 2-minute acquisition post-bleach. Malpighian tubules (MTs) were dissected in Grace’s Insect media with a 1:50 dilution of ProlongLive (antifading agent). A 6% slurry of low-melting agarose (A9414-5G) in Grace’s Insect media was used to stabilize the MTs during imaging on a bridge slide with a cavity to pour the agarose, which was allowed to solidify, forming a soft base to mount the tissue. The intensities recorded in selected regions of interest were obtained using NIS-Elements software. Data fitting and immobile fraction analysis were performed using the NIS-Elements software FRAP analysis module.

### *In-vivo* live imaging

Salivary gland expressing Hrp38GFP (nuclear) in control and *clamp^delPrLD+6Q^*/*clamp^delPrLD^* male and female third instar larvae were dissected in Grace’s Insect media with a 1:50 dilution of ProlongLive (antifading agent) and mounted in the dissecting media on a bridge slide with low-melting agarose (A9414-5G) in Grace’s Insect media as a base. Moving Hrp38GFP condensates in the nucleus were imaged using a 488 nm laser on a Nikon Spinning Disc Confocal Microscope at 60X magnification for 2 minutes each, without any delay in acquisition.

## Computational Methods

### CUT&RUN Data analysis

Sequenced reads were run through FASTQC^56^(fastqc replicate_R1_001.fastq.gz replicate_R2_001.fastq.gz) with default parameters to check the quality of raw sequence data and filter out any sequences flagged for poor quality. Sequences were trimmed and reassessed for quality using TrimGalore (https://github.com/FelixKrueger/TrimGalore/issues/25) and FastQC^56^, respectively. All Illumina lanes of the same flow cell. fastq files were merged, and sequenced reads were mapped to the release 6 *Drosophila melanogaster* genome (dm6). We compared Bowtie2^57^, HISAT2^58^, and BWA^59^. We found the best alignment quality with BWA and thus used its results downstream.

Next, we performed conversion to BAM and sorting (e.g., using: bowtie2 -x dm6_genome -1 replicate_R1_001.fastq.gz -2 replicate_R2_001.fastq.gz -S out. sam> stout.txt 2> alignment_info.txt; samtools view -bSout.sam>out.bam; rm -rfout.sam; samtools sort out.bam -o out.sorted.bam). We removed reads (using samtools) with a MAPQ less than 30 and any PCR duplicates (identified using Picard MarkDuplicates 2.20.2). Peaks identified using MACS2^60^(macs2 callpeak -t out.sorted.bam -B -f BAM --nomodel --SPMR --keep-dup all -g dm --trackline -n outname --cutoff-analysis --call-summits -p 0.01 --outdiroutdir) and keep duplicates separate. To calculate fold-enrichment, macs2 is rerun (macs2 bdgcmp -t $treat -c $control -o $out.sorted.bam_FE.bdg -m FE 2> $ out.sorted.bam_FE.log; macs2 bdgcmp -t $treat -c $control -o $out.sorted.bam_logLR.bdg -m logLR -p 0.00001 2). For motif analysis, the MEME^61^ suite was used. Data was submitted to the GEO repository (#GSE174781, #GSE220981, and #GSE220053).

### iCLIP Data analysis

The method from Heyl et al. 2020^53^ using the Galaxy CLIP-Explorer was followed to preprocess, perform quality control, post-process, and perform peak calling. UMI tools were used for preprocessing, and then UMI tools and Cutadapt were used for Adapter, Barcode, and UMI removal. Cutadapt (Galaxy version 3.5) was used for filtering with a custom adaptersequenceAGATCGGAAGAGCGGTTCAGCAGGAATGCCGAGACCGATCTCGTATGCCGTCTTCTGCTTG. All other settings followed the Heyl et al. 2020 Galaxy iCLIP-explorer workflow. UMI-Tools Extract (Galaxy Version 1.1.2+galaxy2) was then used with a barcode pattern of NNNXXXXNN. No unpaired reads were allowed. The barcode was on the 3’ end. Je-Demultiplex (Galaxy Version 1.2.1) was then used for demultiplexing. FastQC was used for quality control. Mapping was done by RNA STAR (Galaxy version 2.5.2b-2) using dm6. All settings were chosen based on the existing parameters from the iCLIP-explorer settings. We selected FALSE for the option to use end-to-end read alignments with no soft-clipping. bedtools used for Read-Filtering, and UMI-Tools (Galaxy version 0.5.3.0) for de-duplication. PEAKachu was used for Peak Calling to generate bed files. The PEAKachu settings were followed using the Galaxy CLIP-explorer workflow. The maximum insert size was set to 150, the minimum cluster expression fraction was set to 0.01, the minimum block overlap was set to 0.5, and the minimum block expression was set to 0.1. The Mad Multiplier was set to 0.0, the Fold Change Threshold was set to 2.0, and the adjusted p-value threshold was set to 0.05. Peaks were annotated using RCAS^62^ (RNA Centric Annotation System), an R package using RStudio. MEME Suite is used for motif detection. RCAS was used to analyze the functional features of transcriptomes isolated by iCLIP. Data was submitted to the GEO repository (#GSE205987).

### Integrating CUT&RUN and iCLIP data

A Python script was created that iterates through all of the DNA peak bed files for CLAMP DNA binding sites in Kc and S2 cell lines (CUT&RUN data, #GSE220053) as a reference and tests for overlap with CLAMP-bound RNA peaks (each sequence is between 25-50bp in size) in the Kc and S2 (iCLIP data, (#GSE205987). The overlaps are categorized into four main categories based upon the location of the overlap: 1) completely overlapping (purple lines in frequency plot), 2) partially overlapping at the DNA peak start site (red lines in frequency plot); 3) partially overlapping at the DNA peak end site (blue lines in frequency plot) and 4) non-overlapping, i.e. when there is an overlap in a region outside the DNA binding site (yellow lines in frequency plot). This extended region is defined by the *scope* variable in the script, allowing the overlap to look for binding sites near the DNA binding site (this scope is 2 kb, including the DNA binding site). We note that multiple RNA peaks can be found within a single DNA peak. These overlaps are placed onto a [-scope, scope] region. Then, each type of overlap, shown in a different color, is overlaid and plotted on a frequency plot. So, if the frequency at a given base pair is 5, five overlaps contain that base pair within the region defined by the scope.

### Analysis of Imaging data

#### *In vivo* co-localization of CLAMP and Hrp38

Images of the fixed salivary glands immunostained for CLAMP (magenta, anti-CLAMP antibody) and Hrp38 (green, GFP) were acquired as z-stacks and saved as nd2 files. The raw image data from each nd2 file was converted to 3D numpy arrays for each specified channel (magenta and green) employing the nd2 reader Python package. Colocalization analysis was then performed to determine the Pearson Correlation Coefficient (PCC) and corresponding p-value between the two channels of each nd2 file using the scikit-image Python package (scikit-image 0.22.0), indicating the degree of overlap between the proteins (CLAMP and Hrp38). The PCC and p-value were obtained for 20 male controls, 38 male *clamp^delPrLD+6Q^*/*clamp^delPrLD^*, 22 female controls, and 28 female *clamp^delPrLD+6Q^*/*clamp^delPrLD^*nd2 files. The mean PCC and standard error of the mean (calculated using SciPy) for each background across all nd2 files were then plotted using the matplotlib Python package. Statistical analysis was performed to compare colocalization coefficients across different experimental conditions, including pairwise p-values and t-values for each experimental background relative to the others.

All image processing and analysis for colocalization were done through a custom Python script. (https://github.com/smritivaidyanathan/3D_Speckle_Volume_Quantifier/blob/main/scripts/coloc <u>alizer.py</u>).

#### Quantification of speckle dynamics; Bound Fraction calculations

Live-imaging movies (2 min long) of dynamically moving GFP-tagged Hrp38 protein were pre-processed using Fiji (ImageJ) Python macro script, involving Fiji Plugins and built-in software on the Hrp38 condensate (green channel) to distinguish Hrp38 phase condensates from the cell background. Simple Ratio bleach correction was applied, and the minimum and maximum intensity adjustment was executed based on the mean intensity and standard deviation of intensity. The specific adjustment method varied depending on the image quality, ensuring optimal visibility of Hrp38 condensate speckles. Afterward, background subtraction was performed with a rolling ball radius of 120 pixels. The conversion to a binary image involved max entropy thresholding, although the minimum thresholding method was employed for some images to accommodate diverse lighting conditions. The thresholder images were saved as TIFF files for use in the TrackIt tracking software^44^, a program for tracking and analyzing single-molecule experiments developed by the Gebhardt lab. In total, 29 male control movies, 55 male *clamp^delPrLD+6Q^*/*clamp^delPrLD^*mutant movies, 23 female control movies, and 65 female *clamp^delPrLD+6Q^*/*clamp^delPrLD^*mutant movies were utilized in subsequent analyses.

For tracking analysis, TIFF movies from each experimental condition were loaded into TrackIt^44^, and analysis was performed using MATLAB on a Windows 10 PC, utilizing default settings with specific parameter adjustments: threshold (4), tracking radius (8), minimum track length (2), one gap frame allowed, and minimum track length before the gap (3). Each experimental condition was analyzed using TrackIt’s MATLAB-based data visualization tool, with settings further edited for formatting in a MATLAB script. Tracking data outputs were used for bound fraction analysis according to the TrackIt package^44^. TrackIt provided information on tracked events, including long, short, linked, and non-linked events. The calculation of bound fractions by TrackIt involved determining the ratio of the sum of short and long events to non-linked events. Movie-wise bound fraction values and the pooled fraction were plotted (sum over all movies for each type of event). Statistical analysis comparing the bound fraction of Hrp38 speckles across different experimental conditions was performed using a paired Student’s t-test for each experimental background against the others. Error bars for the pooled fraction values were calculated using linear error propagation, following the formula indicated in the TrackIt Manual^44^ (https://gitlab.com/GebhardtLab/TrackIt/-/blob/master/TrackIt_manual.pdf?ref_type=heads).

#### Quantification of speckles volume

Z-stack confocal images from fixed third instar salivary gland tissue of male and female wildtype control and *clamp^delPrLD+6Q^*/*clamp^delPrLD^*mutant samples expressing GFP tagged Hrp38 protein in the nucleus were used to determine the size of Hrp38 nuclear speckles in the presence of CLAMP^WT^ and CLAMP^delPrLD^. We wrote a Python script for the pre-processing step to create a binary image, with all areas covered by a fluorescently tagged Hrp38 condensate to be represented by a 1, and all others to be marked as 0, using the following steps: a. Conversion of the z-stacks to a 3D array using the nd2reader Python package. b. Smoothing the image to reduce noise using a Gaussian filter with a sigma value of 1.0. c. Thresholding the image using the formula, where the threshold was calculated as follows:

Each pixel in the smoothed image is represented by an intensity value ranging from 0 (black) to 255 (white). Following the application of a smoothing algorithm, the arithmetic average (μ) and standard deviation (σ) are calculated across all pixel intensities in the smoothed image.

Thresholding was performed to generate a binary image using these calculated parameters. The threshold value (T) is defined as:

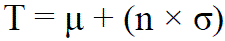

where n is a user-defined constant, with n≥0, for each pixel in the smoothed image, if the intensity was greater than T, the corresponding pixel in the binary image was assigned a value of 1 (white). Conversely, if the intensity was less than or equal to T, the corresponding pixel in the binary image was assigned a value of 0 (black). The resulting binary image retained the same dimensions and pixel count as the original smoothed image.

d. Storing and sending to the next step of the pipeline. In the above function, the value of n varied between datasets. It was refined through a visual inspection of the thresholded 3D image using the Napari Viewer python package (https://napari.org), and n was set to equal to 7. In total, 17 male control, 15 male *clamp^delPrLD+6Q^*/*clamp^delPrLD^*mutant, 17 female control, and 25 female *clamp^delPrLD+6Q^*/*clamp^delPrLD^*mutant z-stack movies were processed and analyzed. To account for irregular shapes of speckles, the segmentation algorithm union-find^63^ with modified connectivity parameter set as None, meaning that the program will not limit the number of jumps, set to input.ndim, or the size of the array itself (Skimage. measure, 2024) was used. The mean speckle size for each category was plotted with the 95% confidence intervals, calculated using a Z-score of 1.96. The same was done for medians, except using bootstrapping with 1000 samples to calculate the error bars, again calculated using a Z-score of 1.96. The pairwise p-values and t-values were calculated using an independent sample t-test (student t-test). Details of the 3D volume quantification scripts can be found on the Github page: https://github.com/smritivaidyanathan/3D_Speckle_Volume_Quantifier.

## Acknowledgments

This work was supported by R35GM126994 to E.N.L. from NIH, and by National Science Foundation (Directorate for Biological Sciences) 1845734 and R01GM147677 to N.L.F. A.M.C is funded by the NSF Graduate Research Fellowship and CCMB, Brown University. We thank Bloomington Stock Center for fly lines. We thank Steve Henikoff for the pAMNase protein and spike-in DNA for Cut and Run.

## Author Contributions

**M.R**., **N. L. F,** and **E.N.L**. planned experiments, analyzed results, and wrote the manuscript. **J.Z** did all protein purifications and the NMR and phase separation assays for CLAMP. **M.R.** conducted the experimental work and collected data for Cut and Run, iCLIP, FRAP, Live imaging, RNA-EMSA, RNA isolation for RNA-sequencing, and generated fly mutants. **P.M** analyzed third instar larval RNA-seq data using the time2splice pipeline to identify CLAMP-dependent splicing events, integrated CLAMP iCLIP data with CLAMP CUT&RUN in cell lines, and performed the splicing assay for *hrp38*. **S.V.** analyzed all the imaging data sets. **R.V.** performed the co-LLPS assay with Hrp38 and CLAMP. **J.S** cloned CLAMP1-300 and CLAMP1-300del PrLD clones. **J.S.**, **V.C., I.P.,** and **K.C.** helped with RNA-EMSA assays. **A.H.** analyzed the iCLIP-seq data. **A.M.C** analyzed CUT&RUN data. **S.H.W.** helped with NMR analysis. **V.J.** and **N.W.** helped with CLAMP Protein Expression and Purification. **A.E.C** and **R.P** helped with the protein and NMR assay.

## Supplementary Figure legends

**Figure S1. A-C** IGV genome browser view showing relative CLAMP DNA and RNA binding at *mrj* (**A),** *sf3b3* **(B),** and *sqd* **(C)** genes.

**Figure S2.** RNA-EMSA showing the binding of increasing amounts of MBP fusion CLAMP-Full length and CLAMP PrLD domain deleted protein to *roX2* RNA biotinylated probes without the CLAMP binding sequence. Concentrations (µM) of CLAMP protein increase from left to right. CLAMP^delPrLD^ and CLAMP^WT^-Full length MBP fusions were used at increasing concentrations of 0, 3.1, 6.2, and 12.5 µM with 200 nM biotinylated RNA probes.

## Table legends

**Table 1a:** CLAMP RNA targets identified in the nuclear fractions of male (S2) and female (Kc) cells.

**Table 1b:** List of genes and genomic locations where CLAMP RNA peaks completely overlap with CLAMP DNA peaks (CRO), CLAMP RNA peaks overlap with 5’ and 3’ ends of DNA peaks (ORF and ORE), and CLAMP RNA peaks are proximal to DNA peaks (PXP) in male (S2) and female (Kc) cells.

**Table 2a:** List of all CLAMP PrLD-dependent differentially spliced genes in *Drosophila* third instar larvae (L3).

**Table 2b:** List of all CLAMP PrLD-dependent differentially sex-specifically spliced genes in *Drosophila* female third instar larvae (L3).

**Table 2c:** List all CLAMP PrLD-dependent differentially sex-specifically spliced genes in *Drosophila* male third instar larvae (L3).

**Table 3:** List of CLAMP PrLD-dependent male and female specifically spliced genes whose RNA isoforms are direct targets of the CLAMP protein.

## Movie legends

**Movie 1:** Live imaging of dynamic Hrp38GFP in the control male third instar larval salivary gland cell nucleus.

**Movie 2:** Live imaging of dynamic Hrp38GFP in *clamp*^delPrLD^/*clamp*^delPrLD+6Q^ male third instar larval salivary gland cell nucleus.

**Movie 3:** Show recovery of Hrp38GFP during FRAP experiment in the control (female) third instar larval Malpighian tubule cell nucleus.

**Movie 4:** Show recovery of Hrp38GFP during FRAP experiment in *clamp*^delPrLD^/*clamp*^delPrLD+6Q^ (female) third instar larval Malpighian tubule cell nucleus.

