## Supplementary figures and images for "A dual DNA/RNA-binding factor regulates co-transcriptional splicing through target RNA interaction and modulates splicing factor dynamics"

### Supplementary Figures

Figure S1 Ray, Zaborowsky et al 2026

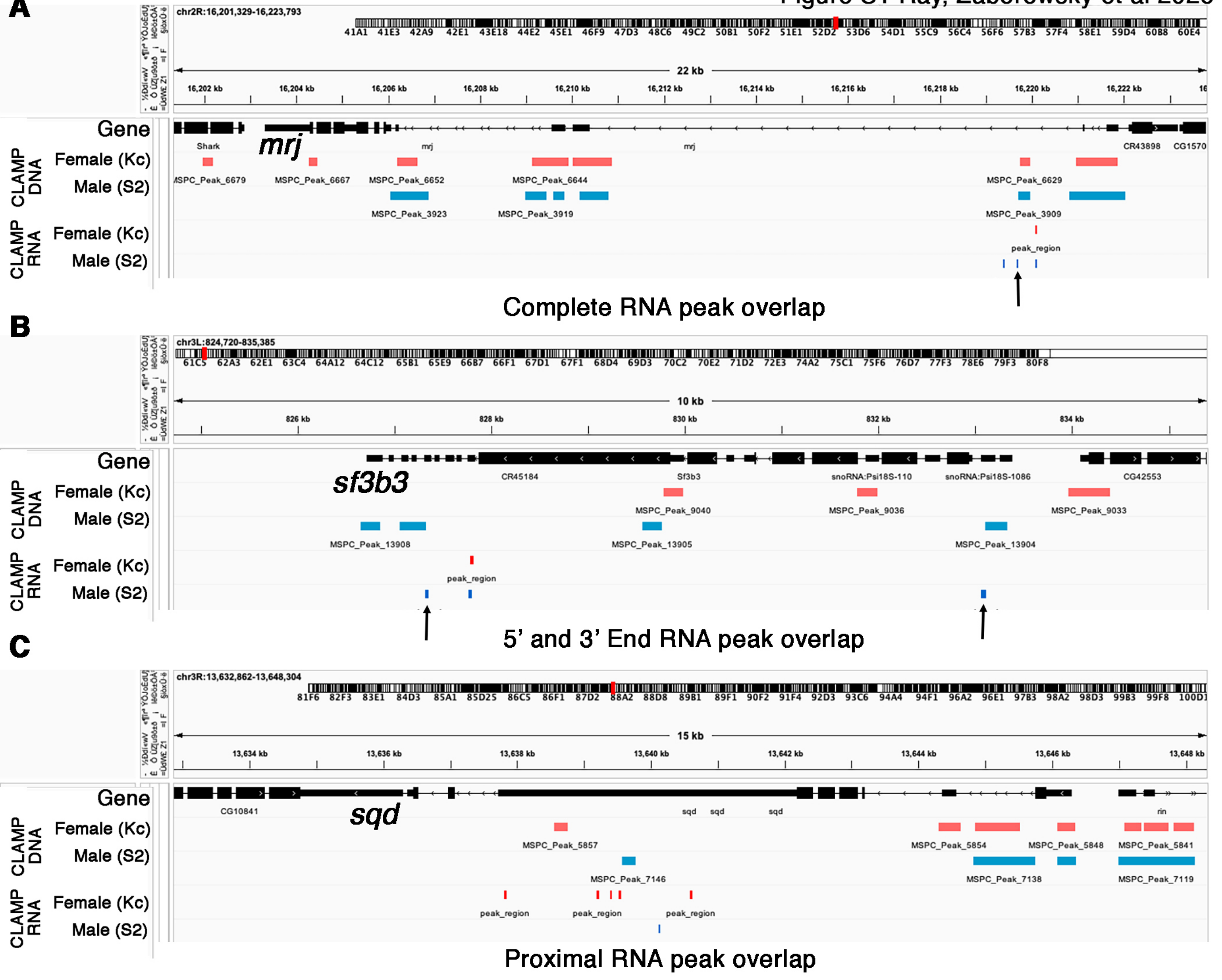

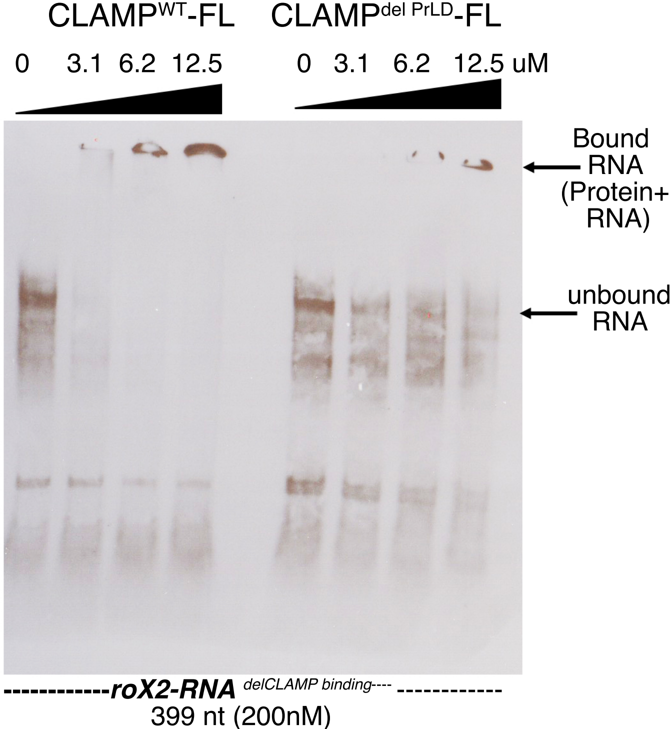
